# Exploring mechanisms of change in a Southern Ocean fishery with a co-produced network model

**DOI:** 10.1101/2024.10.23.619600

**Authors:** Delphi Ward, Nicole Hill, Jessica Melbourne-Thomas, Dirk Welsford, Rhys Arangio, Malcolm McNeill, Simon Wotherspoon, Philippe Ziegler, Stuart Corney

## Abstract

A key challenge in planning long-term fisheries sustainability is overcoming uncertainties in predicted changes in target populations and catch rates in response to climate change and fishery trends. We combine transdisciplinary knowledge co-production and qualitative network modelling to advance system understanding and elucidate likely responses of a Patagonian toothfish fishery to future change. We co-developed a model of the Kerguelen Plateau biophysical-socioeconomic system with knowledge holders from industry and science; the first whole-of-system qualitative network model of intermediate complexity for this region. We present new approaches for dealing with uncertainty in network structure, and for investigating how perturbations propagate through a network. Using these tools, we found multiple potential pathways of decline for toothfish population and catch, but also some possible mechanisms of increase dependent on magnitude of certain effects. Our results highlight critical information gaps, including the potential role of scavenging benthos in fishery-ecosystem interactions and likely changes in the prey-field in response to warmer water, that require filling to improve predictions for toothfish populations and catch and forward planning for the fishery.

## Introduction

Interacting climate change impacts on ocean environments and biota across scales are creating conditions beyond what marine ecosystems have previously experienced, are increasing the challenge of sustainable management (Lotze et al. 2019; Bindoff et al. 2019; Cheung et al. 2021). Managing fisheries sustainably into the future requires predicting how fish stocks will be affected by environmental and ecological changes as well as how fisheries behaviour will interact with these changes (Holsman et al. 2020; Galappaththi et al. 2022). Doing so for deep and remote areas is particularly challenging with limited data about community structure, dynamics and responses to change. Addressing these challenges effectively calls for interdisciplinary collaborative efforts to proactively explore future needs and ecosystem trajectories (Wyborn et al. 2019).

Joint problem-framing and knowledge co-production among stakeholders are recognised as effective means of addressing these broad challenges (Norström et al. 2020; Pearce and Ejderyan 2020). This represents an interactive approach to science delivery to ensure the needs of all stakeholders are addressed, and facilitate a shift towards ecosystem-based (as opposed to stock) management. An important step in this process is to develop a shared understanding, framework and language for the system, desired outcomes and knowledge gaps that can be addressed through research. This can be achieved through collaborative development of a conceptual model of the system, including its important features and processes (Margoluis et al. 2009), and by constructing it together, e.g. on a whiteboard, each participant can see their contributions and how they fit into the larger picture. The benefit of this approach is that it allows diverse stakeholders to be heard at the table and thus achieves both expert knowledge in the findings, and also a sense of ownership. The conceptual model can then be translated into a qualitative network model (QNM) to enable formal testing and exploration to determine likely qualitative responses to sustained (press) perturbations (e.g. Dambacher et al. 2007; Melbourne-Thomas et al. 2013; Reum et al. 2020).

Qualitative network models (QNMs) are based on the sign and direction of interactions between variables, but allow their shape and strength (i.e. mathematical formulation) to remain undefined (e.g. Puccia and Levins 1991; Dambacher et al. 2002; Dambacher et al. 2009). They are particularly useful for exploring systems where interactions are unquantified. They are also relatively quick to construct, meaning that multiple model versions can be created to explore any uncertainties in network structure. Here we use simulation-based QNMs to explore change in the Patagonian toothfish population, fishery and associated ecosystem around Heard Island and McDonald Islands (HIMI) in the Indian Ocean sector of the Southern Ocean. In doing so, we develop and present a new approach for assessing the relative importance of different mechanisms of change within a network.

Patagonian toothfish (Dissostichus eleginoides) are a large, long-lived bentho-pelagic nototheniid fish. They can live for 50 years or more, reaching lengths greater than 2.3 m and weights over 200 kg (Collins et al. 2010). As they mature, toothfish gradually migrate down slope to inhabit deeper waters (>2000 m), and their diet concurrently shifts from being mostly piscivorous as juveniles to become more opportunistic and including more carrion, cephalopods and crustaceans as adults (Collins et al. 2010; Péron et al. 2016). Similarly, toothfish predators change with age; larval and juvenile toothfish are predated by shallow-diving predators, such as larger fish, penguins and fur seals, while larger adults are predated by deep-diving species including sperm whales, large squids and southern elephant seals (Collins et al. 2010). Patagonian toothfish are a high-value species, with medium to large individuals fetching higher market prices. Therefore, fisheries concentrate effort in deeper water (>400m) on the shelf slope where these individuals reside (Péron et al. 2016).

The HIMI Patagonian toothfish longline fishery operates within the Australian Exclusive Economic Zone on the northern Kerguelen Plateau. It is managed by the Australian Fisheries Management Authority (AFMA) in accordance with the ecosystem-based management principles of the Commission for the Conservation of Antarctic Marine Living Resources (CCAMLR). The fishery is widely regarded as being sustainably managed, with quotas set following a precautionary approach and stock assessments updated regularly (≤2yrs) based on scientific trawl survey, catch-at-age and tag-recapture data (Brooks et al. 2019; Ziegler and Welsford 2019; MSC 2020; Subramaniam, Melbourne-Thomas, et al. 2020). While quotas and catch are well within sustainable levels, recent variability in catch rates has raised questions as to how toothfish populations and behaviour will respond to future climate-driven environmental change. Over a period of a few months in 2016, there was a sharp decline and subsequent strong fluctuations in fishery catch rates of this species, coinciding with a surface marine heatwave in the area (Blunden and Arndt 2017; Su et al. 2021). The rapid nature of the catch rate fluctuations and subsequent recovery suggested they were not due to a fishery-related decline in toothfish stock. This unexpected behaviour poses challenges for the fishery, highlighting the need to better understand how the fishery, its indicators and long-term viability may be affected by future changes in the system.

We aimed to co-develop a model of the system (combining scientific, management and industry expert knowledge) and then to use this model to explore mechanisms by which environmental, ecological or fishery behavioural changes could affect catch rates and toothfish stocks in the HIMI fishing area.

## Methods

### Qualitative Network Models

Qualitative network models take the form of a signed, directed graph (digraph) of interacting entities (nodes) and the interactions between them (edges). The network captures the sign and direction of effects, but does not require knowledge of effect strengths, making them attractive for use in systems where effects are unquantified (Levins 1974). In a QNM, edges represent per capita effects on nodes that could be described as a set of ordinary differential equations (Levins 1974). QNMs are neither spatially nor temporally explicit; however, they are constructed to represent a defined spatial and temporal domain.

Analysis of QNMs is based on the community matrix (A), from which considerable information can be derived, including qualitative responses to change. Qualitative network models represent a system at equilibrium, but a sustained increase in a node (press perturbation) will cause increases or decreases in other nodes based on the sign and direction of edges, and the effect-chains and feedbacks they collectively form (see supplementary methods for detail). Specifically, elements of the negative inverse community matrix (-A^-1^) indicate how abundances or densities of each node would differ in a new equilibrium state, compared to the initial equilibrium state, following a press increase of another node (i.e. they do not indicate trends) (Nakajima 1992; Dambacher et al. 2002).

For simulation-based QNMs, numerous quantitative community matrices with different interaction strengths are simulated from the qualitative community matrix (e.g. Dambacher et al. 2003; Raymond et al. 2011; Melbourne-Thomas et al. 2012; Ward et al. 2022). Our QNMs were first created in the drawing software Dia (http://dia-installer.de) then implemented in the R-package QPress (Melbourne-Thomas et al. 2012; Wotherspoon et al. 2016), with interaction strengths sampled from a uniform distribution (i.e. between −1 and −0.01 for negative edges or between 0.01 and 1 for positive edges).

### Model construction and validation

Model development followed an iterative process of collaborative workshops and model refinement. Workshop participants included fisheries scientists, fishers, fishery managers, oceanographers, marine biologists and ecological modellers. The initial conceptual model included detailed interactions between fishery behaviours and metrics, environmental and ecological drivers and toothfish populations and behaviour. We then synthesised this information according to the identified drivers of change and mechanisms explaining how perturbations might flow through the interconnected system to affect stock and catch rates (Table 1). These mechanisms were used to ensure the processes and system components considered to be important were captured in subsequent versions of the model, and the drivers were used to establish the test scenarios. The conceptual model was then translated by the research team into a simplified QNM which was iteratively refined during smaller, targeted workshops and through testing and evaluation (see supplementary text for details). Key assumptions embedded in the model are described in the supplementary text.

**Table 1.**
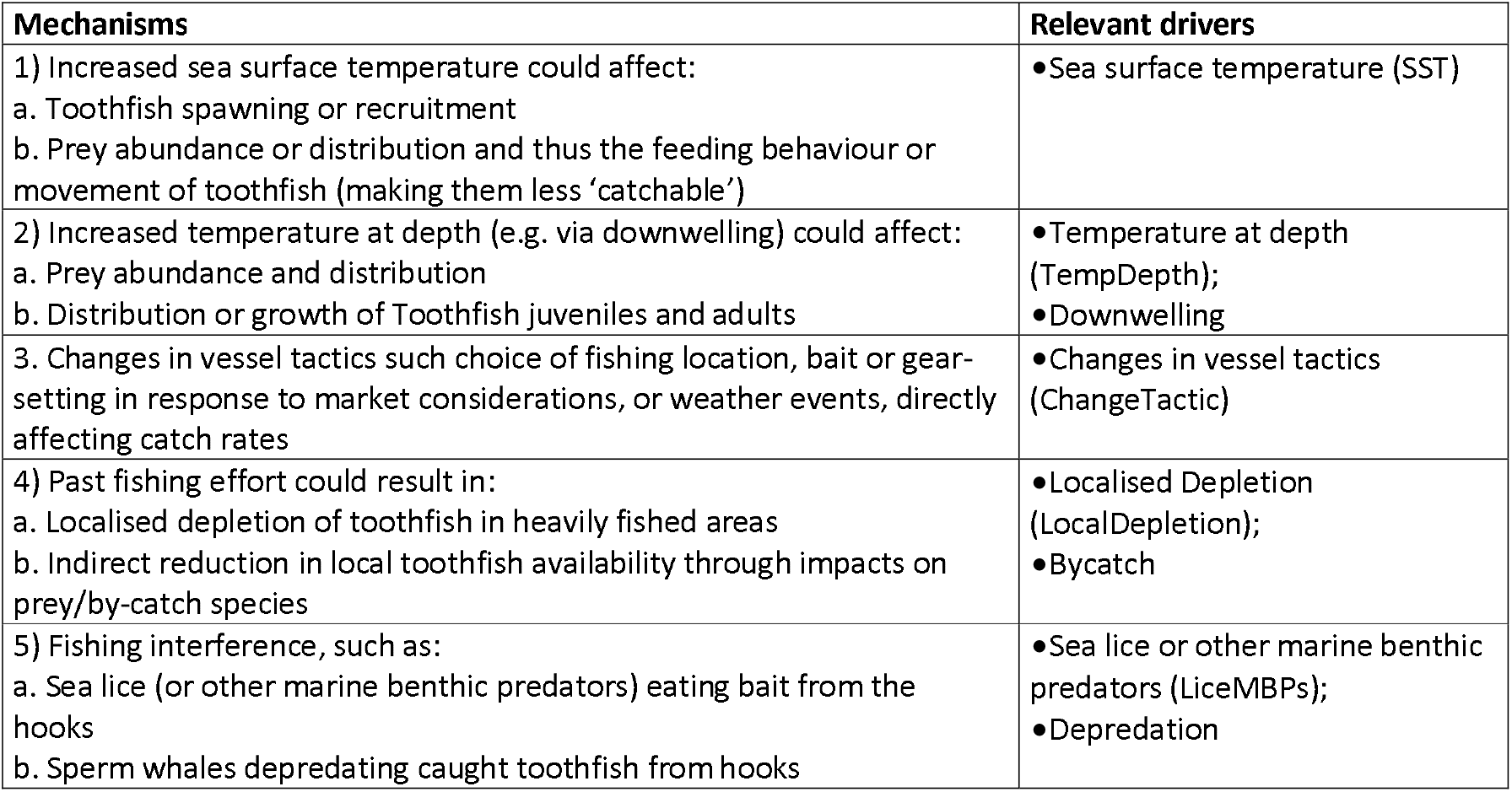
Mechanisms for changes to stock and catch rates (and their associated drivers) derived from a co-produced conceptual model of interactions between fishery behaviours and measures, environmental and ecological drivers and toothfish populations and behaviour. Names of relevant model nodes are given in parentheses beside the driver (when it is an abbreviation).

The few studies that have modelled how different species are likely to respond to increasing water temperatures in the region show complex responses (Subramaniam, Melbourne-Thomas, et al. 2020). Consequently, though our model assumes increased temperature will have some effect on toothfish (big and small), their prey (deep and shallow) and sea lice (or other benthic predators), we do not know the likely sign of these effects. For example, increased water temperature may cause changes in species composition of the toothfish prey field, but whether this translates to an increase or decrease in abundance, accessibility or quality of prey is unknown. Similarly, changing fishing tactic is assumed to affect effort and catch, but the sign of these effects could vary. We captured these alternative possible effect signs in different versions of the model, such that each of the 128 model versions contain a unique combination of these alternative effects. For each model version, 10,000 quantitative community matrices were simulated, where each simulated matrix has a different combination of quantitative edge weights, as described above.

The final model is shown in Figure 1, and summary descriptions of each node and their associated edges can be found in the supplementary material (STable S1). Note that where we refer to a specific model node we use the (capitalised) label corresponding with Figure 1 (e.g. the downwelling node is referred to as ‘Downwelling’).

**Figure 1:**
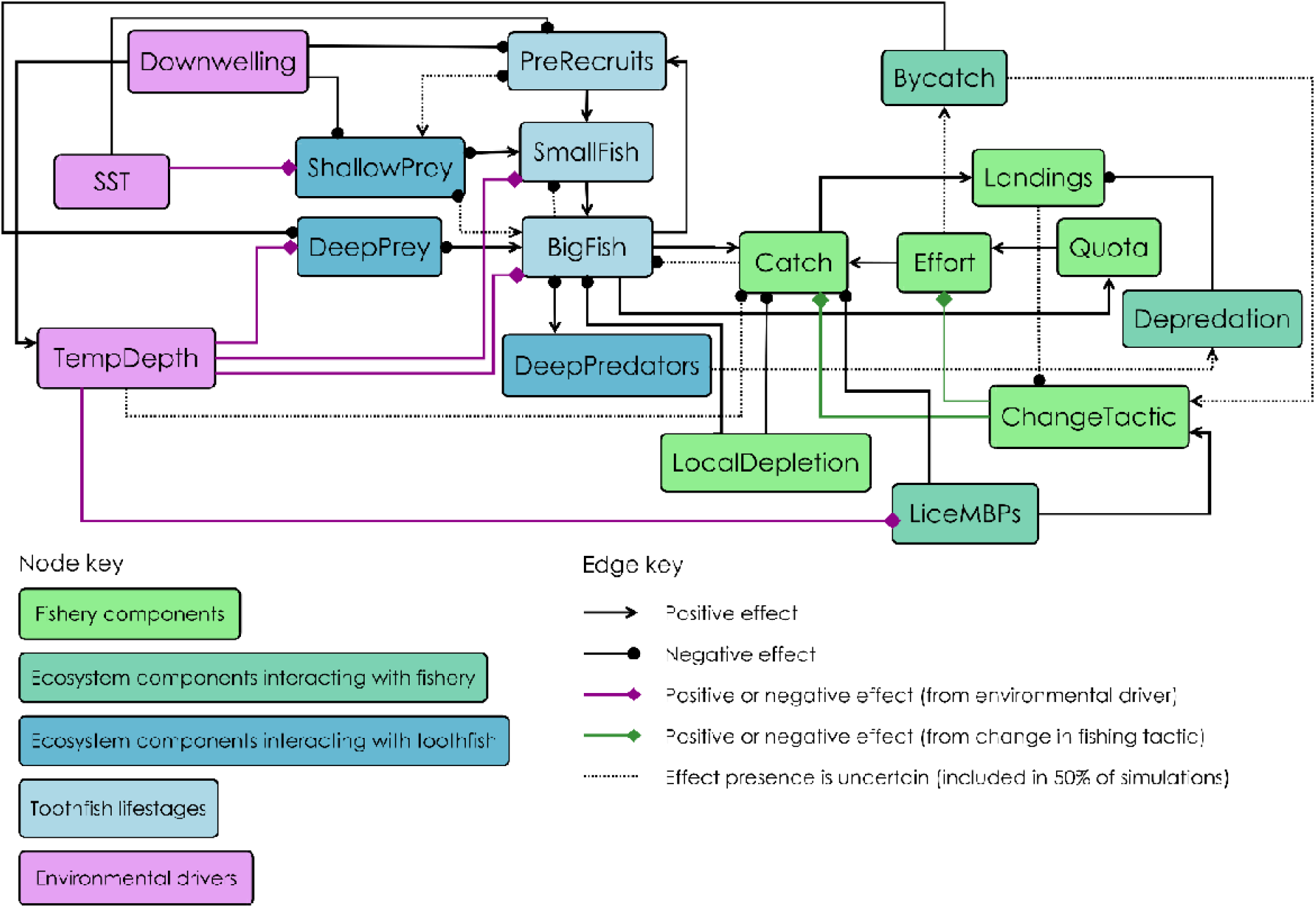
Final qualitative network model of the Heard Island and MacDonald Islands Patagonian toothfish fishery, fish stock, associated ecosystem, and environmental drivers. Three toothfish life stages are shown in pale blue: pre-recruits (representing spawning, egg and larval life stages), small fish (representing toothfish too shallow to be targeted by the fishery) and big fish that are targeted by the fishery. Darker blue nodes represent prey and predators of toothfish at different life stages. Fisheries catch and behaviour nodes are light green. Dark green nodes represent components of the ecosystem which directly affect fisheries: Lice and other marine benthic predators (LiceMBPs), bycatch and depredation. Purple nodes are environmental drivers. Edges terminating in circles represent negative effects, and arrows positive effects, from one node to another. Edges terminating in diamonds indicate effects where the sign of the effect is unknown, and these are either coloured purple (temperature effects of unknown sign) or green (unknown sign of the effects of fisher behaviour – change in fishing tactics). Uncertain presence of an effect is indicated with dotted edges. All nodes have negative self-effects (which are not shown here for clarity).

### Exploring responses to test scenarios (press perturbations)

Based on the set of mechanisms and drivers, eight test scenarios were applied to each of the 10,000 simulated community matrices of each alternative version of the model. Each test scenario involved a sustained (press) increase in one of the following nodes: sea surface temperature (SST), temperature at depth (TempDepth), Downwelling, change fishing tactic (ChangeTactic), localised depletion of toothfish by fishery (LocalDepletion), Bycatch, sea lice or other benthic predators (LiceMBPs), depredation of toothfish from hooks (Depredation). Table 1 shows how these drivers relate to proposed mechanisms. For each node there were a total of 1,280,000 simulated responses to each perturbation, capturing impacts of variation in edge strengths and different combinations of alternative sign effects in determining node response. These responses to perturbation were then analysed to identify 1) mean qualitative responses, 2) the dependence of qualitative responses on unknown effect signs, and 3) relationships between responses and the strength of perturbation effects, as detailed below.

Firstly, responses of each node to each perturbation were averaged across all model versions (including models with positive and negative representations of those edges of unknown sign). Mean responses were scaled between −1 and 1. Average responses were classed high confidence (absolute values between |0.8 and 1|), low confidence (absolute values between |0.4 and 0.8|) or ambiguous (values between −0.4 and 0.4).

Secondly, we wanted to know how key nodes would change in response to perturbation if we made assumptions about the sign of temperature effects and the effects of changing fishing tactic (for cases where the sign of those effects is currently unknown). For each of the 7 effects with unknown sign, half of the model versions represent it as a positive effect and half as a negative effect. For each of these effects we therefore separated the model versions based on the sign of that effect, and compared the mean change in BigFish, Effort, Catch and Landings between the two sets of models. In most cases we did not subset the data further, except for TempDepth and Downwelling perturbations where node responses were sensitive to the sign of effects on multiple nodes. For those two perturbations we identified the two edges for which changing the sign of effects had greatest impact on key node responses (BigFish, Catch and Landings), and present how responses of those nodes differed with different sign combinations of those two effects.

Thirdly, we developed a new approach to identify mechanisms by which perturbations flow through the system, by harnessing information from individual simulations (in contrast to the analyses above in which we worked with mean responses across simulations for each model). The idea is similar to the approach developed by Ward et al. (2022), who examined the relationship between edge strength and network stability to characterise stability conditions. Instead here, we examine the relationship between the strength of perturbation effects (originating from the perturbed node), and the qualitative responses of other nodes. To do this, first, positive responses were assigned the value +1 (increase), negative responses the value −1 (decrease) and responses between 0.001 and −0.001 were assigned the value 0 (no change). For each perturbation, we used the ‘geom_smooth’ function from ggplot2 (Wickham 2016) to plot gaussian GAM smooths of the responses of different nodes against the strength of each of the outgoing effects of the node being perturbed. Cubic splines were used as the smooth type, following the formula y ~ s(x, bs = cs). Sensitivity to effect strengths was assessed based on the shape of the relationships, with greater slope indicating greater dependence of the qualitative response on the strength of an edge. Additionally, for each effect strength to which BigFish and Catch responses were sensitive, the effect strength-response relationships were categorised according to the qualitative response (increase, ambiguous, decrease) with weak and strong effects.

## Results

### Responses of the ecosystem to environmental, ecological or fishery changes

When averaged across all 128 different model versions (with different combinations of effects of increased temperature and changing tactic), some perturbations produced consistent responses (Fig. 2). Sustained increases in SST, localised depletion or bycatch all caused toothfish and catch to decline, and similarly, increased depredation caused a decline in landings. When averaged across model versions, responses of nodes to other press perturbations were ambiguous, either because the models predicted little change, or because the effects of the perturbations differed between model versions (i.e. responses conditional on particular combinations of effect signs). Ambiguity of responses can arise where no change is predicted (e.g. when the node is weakly connected in the model), where positive and negative feedbacks within the network counterbalance each other to create uncertainty in response, or where differences in model versions create bimodal response distributions such that the mean response is close to zero (in other words, it is either the system itself that produces the ambiguity, or, lack of clarity in our understanding of it that produces ambiguity, or a combination of these).

**Figure 2:**
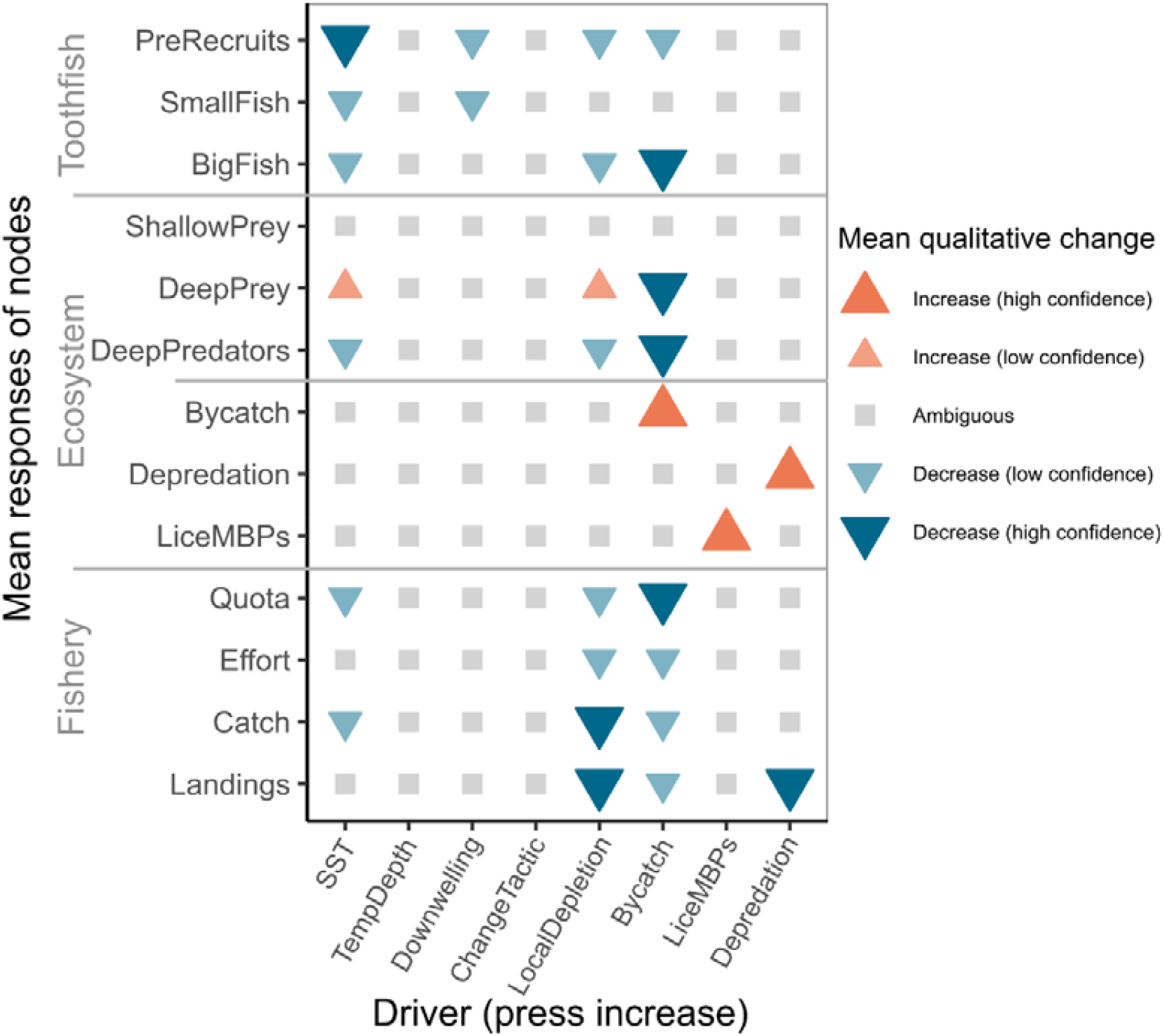
Node responses (along y-axis) to positive press perturbations of driver (along x-axis), averaged across all simulations and all model versions. All responses are scaled between −1 and 1. Ambiguous responses are those with mean values between −0.4 and 0.4. Responses with high confidence are those with mean values between −0.8 and −1 or 0.8 and 1. Node responses (positive or negative) with high confidence indicate the response occurs in most simulations and for most model versions. See Figure 3 for detail regarding the variability in responses among model versions.

### Conditional responses

Figure 3 shows how responses of key nodes for each test scenario are sensitive to edges with unknown sign. Unsurprisingly, responses to environmental perturbations depended on how increased temperature affected different components of the system, and responses to fishery perturbations tended to be conditional on how changes in fishery behaviour (ChangeTactic) affected Effort and Catch. Responses to increased SST depended on the effect of SST on shallow prey: toothfish and catch declined in model versions where abundance and/or quality of prey was negatively affected by higher SST, whereas responses were ambiguous in versions where shallow prey was positively affected by higher SST. Responses to increased TempDepth and Downwelling varied depending on combinations of effect signs. For these test scenarios, large toothfish (BigFish) were most likely to decrease if temperature negatively affected both DeepPrey and BigFish, while Catch was most likely to decrease if temperature positively affected LiceMBPs and negatively affected DeepPrey (Figure 4 and Supp FigS1). Note that in Figure 3, for the LiceMBPs perturbation, the apparent sensitivity of catch response to the effect of TempDepth on BigFish required further investigation as there was no increase in temperature considered in the scenario, and we know no mechanism by which this effect could operate. The difference in means between the sets of simulations was marginal and close to the cut-off between ambiguous vs negative responses. We therefore have low confidence in this result and consider it an artefact. Similarly, for the SST perturbation, the apparent sensitivity of BigFish and Catch responses to ChangeTactic effects appears to be marginal – all the variability in response category is due to the effect of SST on ShallowPrey, but within the set of model versions where SST negatively affect shallow prey, the mean responses of Catch and BigFish were slightly more negative (closer to −1) in model versions where ChangeTactic positively affected Effort and Catch.

**Figure 3:**
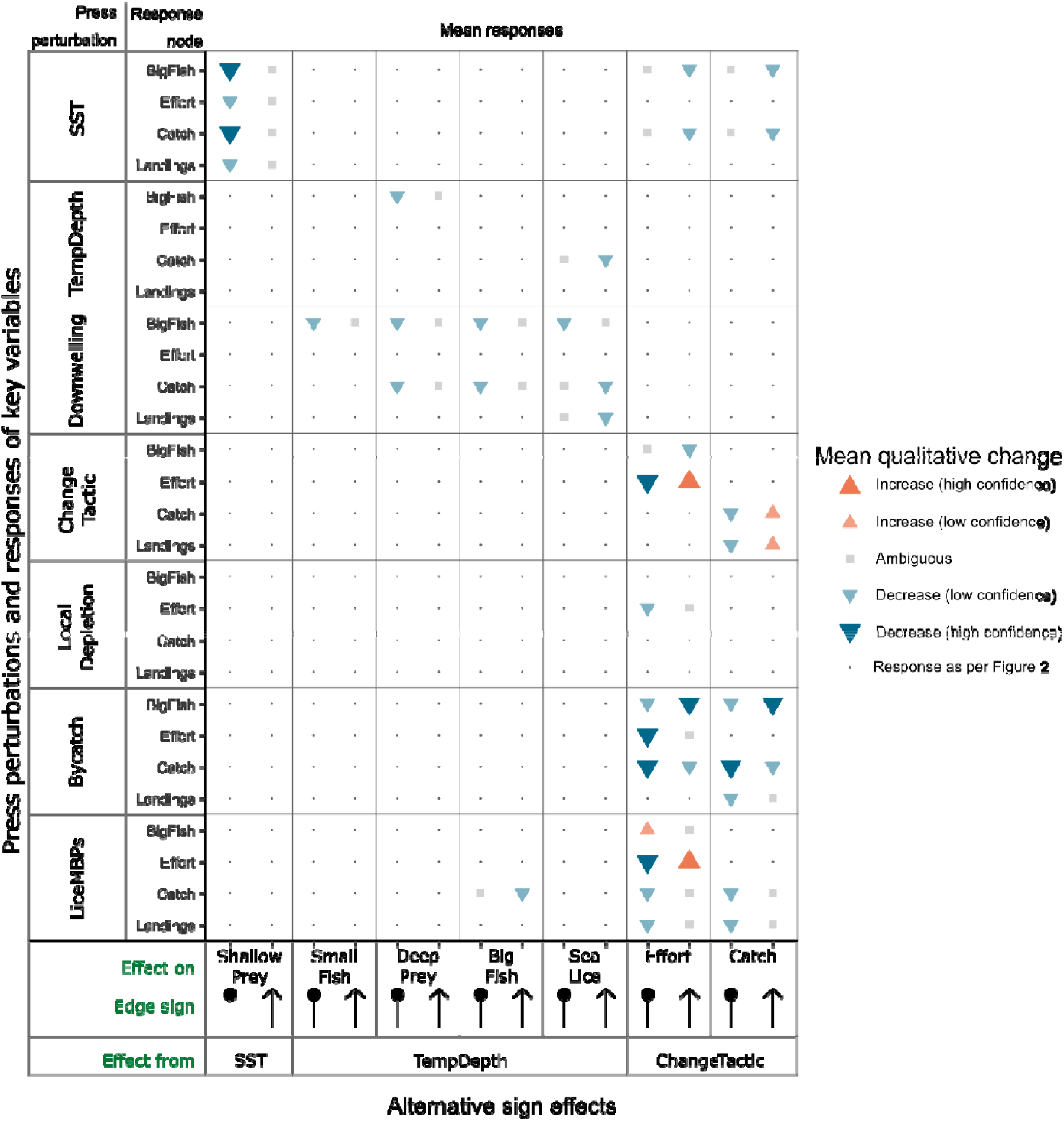
Comparison of mean responses between model versions according to sign of interactions. Each point represents the mean response of a node to a perturbation, for the n=64 (of 128 model versions) in which the edge indicated along the x-axis is either negative (dots) or positive (arrows). The y-axis shows the responses of 4 key nodes (BigFish, Effort, Catch, Landings) to each of the press perturbations (except Downwelling, for which node responses were the same as shown in Figure 2). Each horizontal pair of points within a gridcell thus together represent all 128 model versions, differentiated by the sign of the edge indicated on the x-axis for that gridcell. For clarity, we show only those cases where the mean response differs qualitatively between model sets where the edge is positive vs negative. Points filled with a small dot indicate that there was no discernible difference between the responses of the node in the two sets of model versions. Responses to increased depredation are not shown because they were not sensitive to any of the alternative sign effects (responses as per Fig. 2). All responses are scaled between −1 and 1. Ambiguous responses are those with mean values between −0.4 and 0.4. Responses with high confidence are those with mean values between −0.8 and −1 or 0.8 and 1.

**Figure 4:**
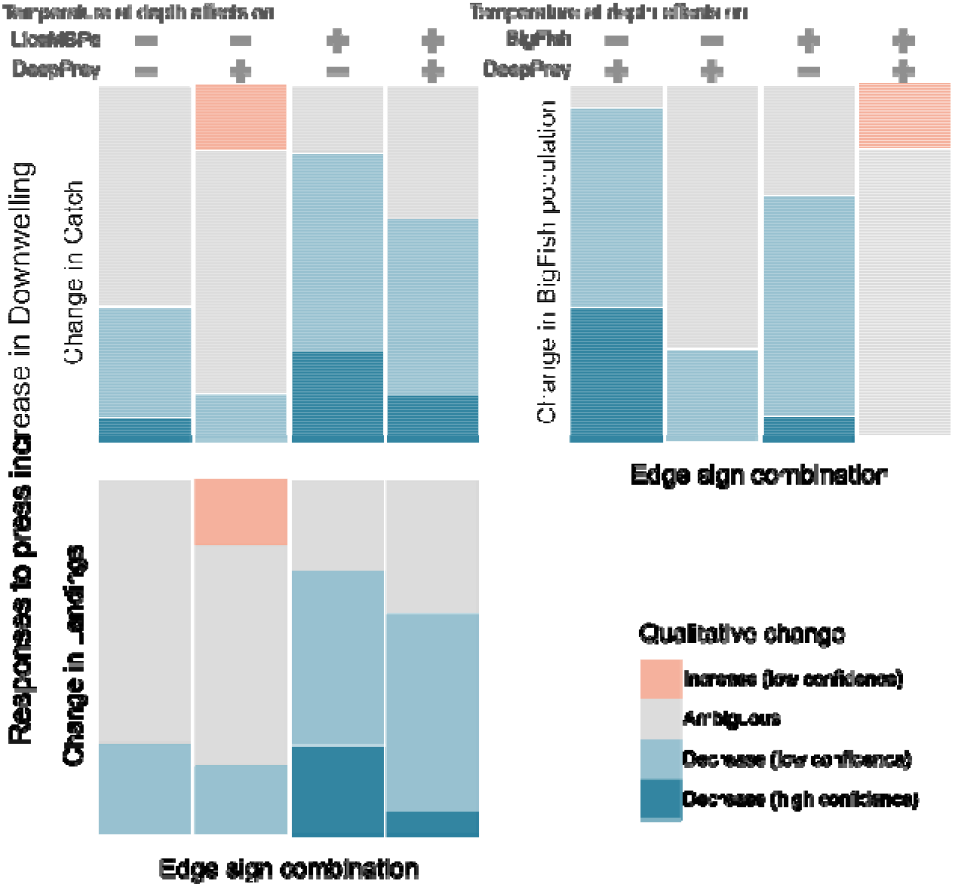
Mosaic plots illustrating how responses to Downwelling press perturbation vary according to sign of temperature effects. Plots in the left-hand column display responses of Catch (upper) and Landings (lower) in model versions grouped by the sign of the effects of TempDepth on Lice and other marine benthic predators, and on Deep Prey of toothfish (the sign of these two edges explain the most variability in responses of Catch and Landings). The plot on the right shows BigFish responses in models grouped by the sign of TempDepth effect on BigFish and on Deep Prey (the sign of these two edges explain the most variability in BigFish response). Each mosaic plot represents the responses of nodes across all 128 model versions, with each column representing the responses in a different subset (n=32) of those.

### Relationships between effect strength and responses to perturbation (mechanisms of change)

Figure 5 shows how the likelihood of qualitative responses (decrease, no change or increase) vary according to the strength of outbound effects from the perturbed node. In some cases, and for some nodes, the strength of an edge affects the likely sign of the response. For example, when increased SST had a weak positive effect on ShallowPrey, then BigFish, Catch and Landings tended to decrease, but they tended to increase when that edge was strong (Fig. 5a, solid yellow, pink and dark-pink lines). In other cases, the likelihood of a decrease (or increase) changed with effect strength. For example, when local depletion weakly affected BigFish (edge strength close to 0), then PreRecruit and BigFish responses were slightly negative (no change or slight chance of decrease), but declines became more likely (responses approaching −1) in simulations where the effect was stronger (Fig. 5d). Similarly, Landings and Catch responded ambiguously to increased LiceMBPs when the effect of lice on Catch was weak, but tended to decline when the effect was strong (Fig. 5b, dark-pink and light-pink dashed lines). In other cases, the strength of an effect did not modulate the likelihood of a qualitative response. For example, Landings consistently declined in response to increased depredation whatever the strength of that effect (Fig. 5f, dark-pink line).

**Figure 5:**
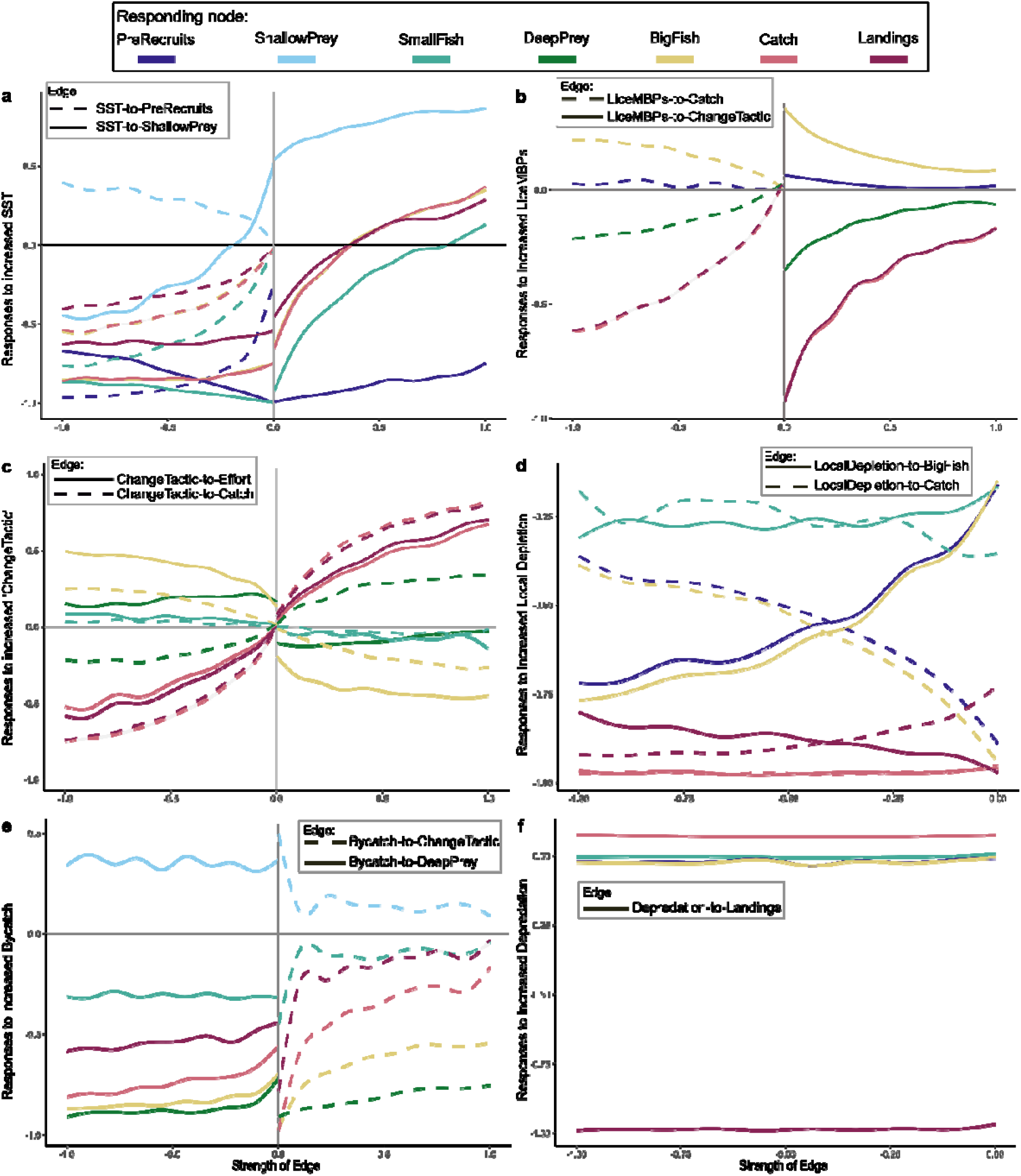
Relationships between qualitative responses to perturbation and effect strengths. Each plot shows responses to one of the press perturbations: a) higher SST, b) more sea lice or other marine benthic predators, c) increase in ChangeTactic, d) more LocalDepletion, e) higher Bycatch and f) higher Depredation. Each line is a gam smooth of the qualitative responses of one node against one of the outbound edges of the node being perturbed. Colours correspond to the response of one node to that perturbation (note: for clarity, not all nodes are shown in each subplot). Line styles correspond to a particular effect from the node being perturbed. Values on the x-axis correspond to the strength of the outward effects from the node being perturbed. Values on the y-axis correspond to the qualitative responses of the nodes, with −1 indicating a decline, +1 an increase, and 0 indicates no change (values between |0 and 1| suggest a mix of qualitative responses). The 95% confidence intervals are plotted in grey but are too narrow to distinguish from the lines. Equivalent plot for downwelling perturbation shown in Supp. Fig. S2.

Fisheries behaviour affected responses to perturbation. When there was a press increase of ChangeTactic, the response of Catch was most sensitive to the strength of the effect on Effort and BigFish response was more sensitive to the strength of the effect on Catch (Fig. 5c, pink and yellow lines). However, the behavioural response of the fishery to other changes within the system are also important for how toothfish stock and catch change. When there was a press increase in Bycatch, then Catch and Landings consistently declined no matter the strength of the effect of Bycatch on DeepPrey (Fig. 5e, solid pink and dark-pink lines). However, Catch and Landings responses were more sensitive to the strength of the effect of Bycatch on ChangeTactic, that is, Catch and Landings tended to decline when that effect was weak and have ambiguous responses when it was strong (Fig. 5e, dashed light-pink and and dark-pink lines). Similarly, Catch and Landings responses were sensitive the strength of effect of LiceMBPs on ChangeTactic, when there was a press increase in LiceMBPs (Fig. 5b, dashed pink and dark-pink lines).

The relative importance of the various TempDepth effects differed for different components of the system (Figure 6 and Supp FigS3). For example, BigFish and PreRecruit qualitative responses to an increase in temperature at depth varied most according to the strength of the TempDepth effect on DeepPrey (Fig 6a, e, dash-dot blue line), whereas Catch and Landings responses varied most according to the strength of the effect of TempDepth on LiceMBPs (Fig. 6b, d, solid yellow line). In this latter case, most of the variability in Landings response occurred when the effect of TempDepth on LiceMBPs was negative, and the likelihood of decline was consistent when this effect was positive. SmallFish qualitative response was most sensitive to the strength of the direct effect from TempDepth (Fig. 6c), and this effect flowed through the network, but had relatively minor importance for BigFish, Catch and Landings outcomes (Fig. 6a, b, d).

**Figure 6:**
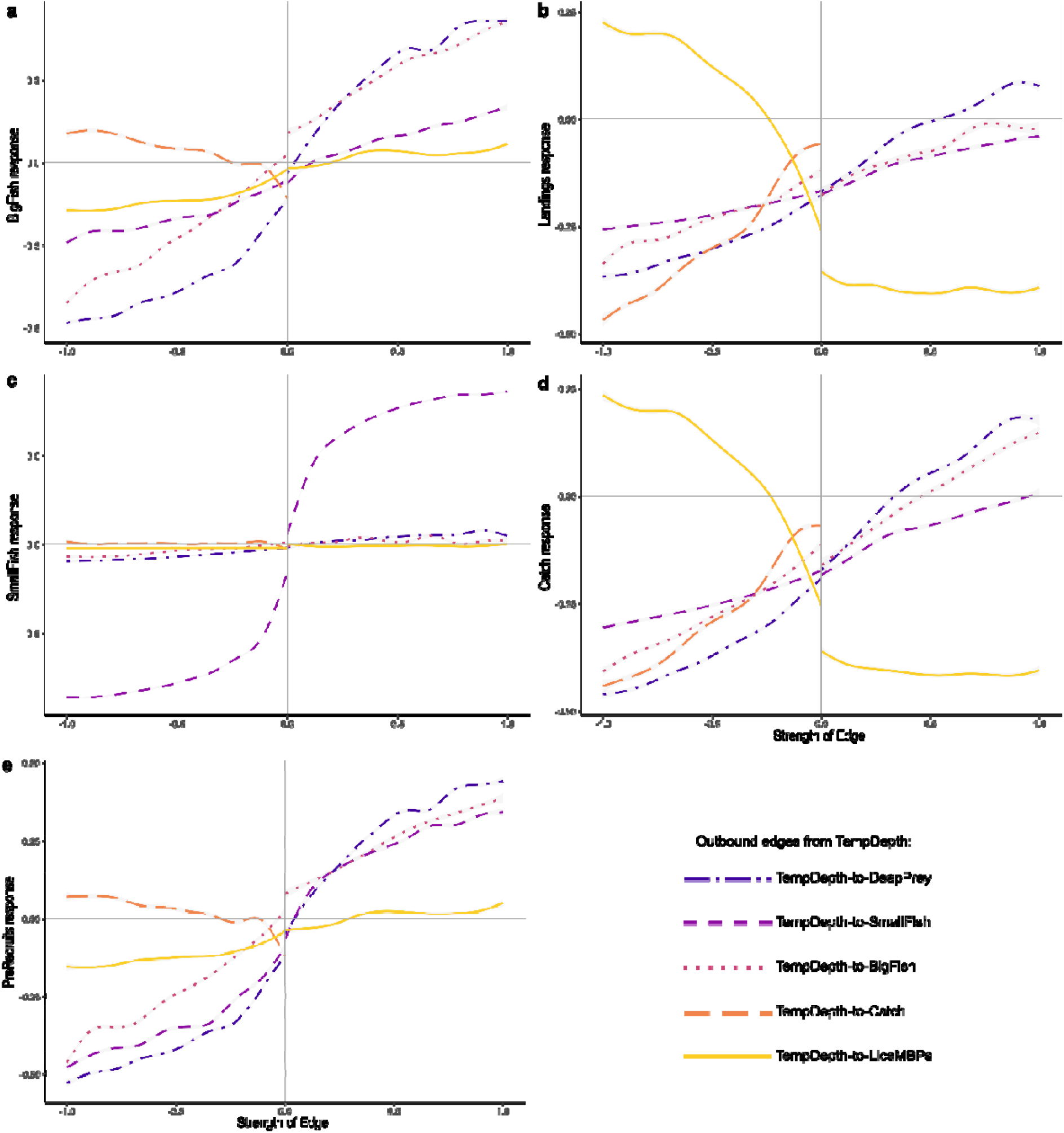
Relationships between qualitative responses to TempDepth perturbation and effect strengths. Each plot shows the responses of one node: a) BigFish, b) Landings, c) SmallFish, d) Catch, and e) PreRecruits. Each line is a gam smooth of qualitative responses of that node one against one of the outbound edges from TempDepth, as indicated by line colour and style. Values on the x-axis correspond to the strength of the outbound edges. Values on the y-axis correspond to the qualitative responses of the nodes, with −1 indicating a decline, +1 an increase, and 0 indicates no change (values between |0 and 1| suggest a mix of qualitative responses). The 95% confidence intervals are shaded grey. Equivalent plots of prey node responses shown in Supp. Fig. S3.

Overall, we observed four categories of strength-response relationships among the edge strengths to which BigFish and Catch responses were sensitive, as described above and summarised in Table 2. The most frequently observed category of relationship was an ambiguous response to perturbation when an edge was weak, and decline when the edge was strong (Table 2, row 1). However, for those edges where the sign is unknown (i.e. effects of temperature and changing fishing tactic), it is possible for BigFish and Catch to increase, if the effects are strong enough and of the right sign (Table 2, rows 3 and 4). In some cases, BigFish and Catch were more likely to decline if an effect was weak, and have an ambiguous response when it was strong (Table 2, row 2).

**Table 2:**
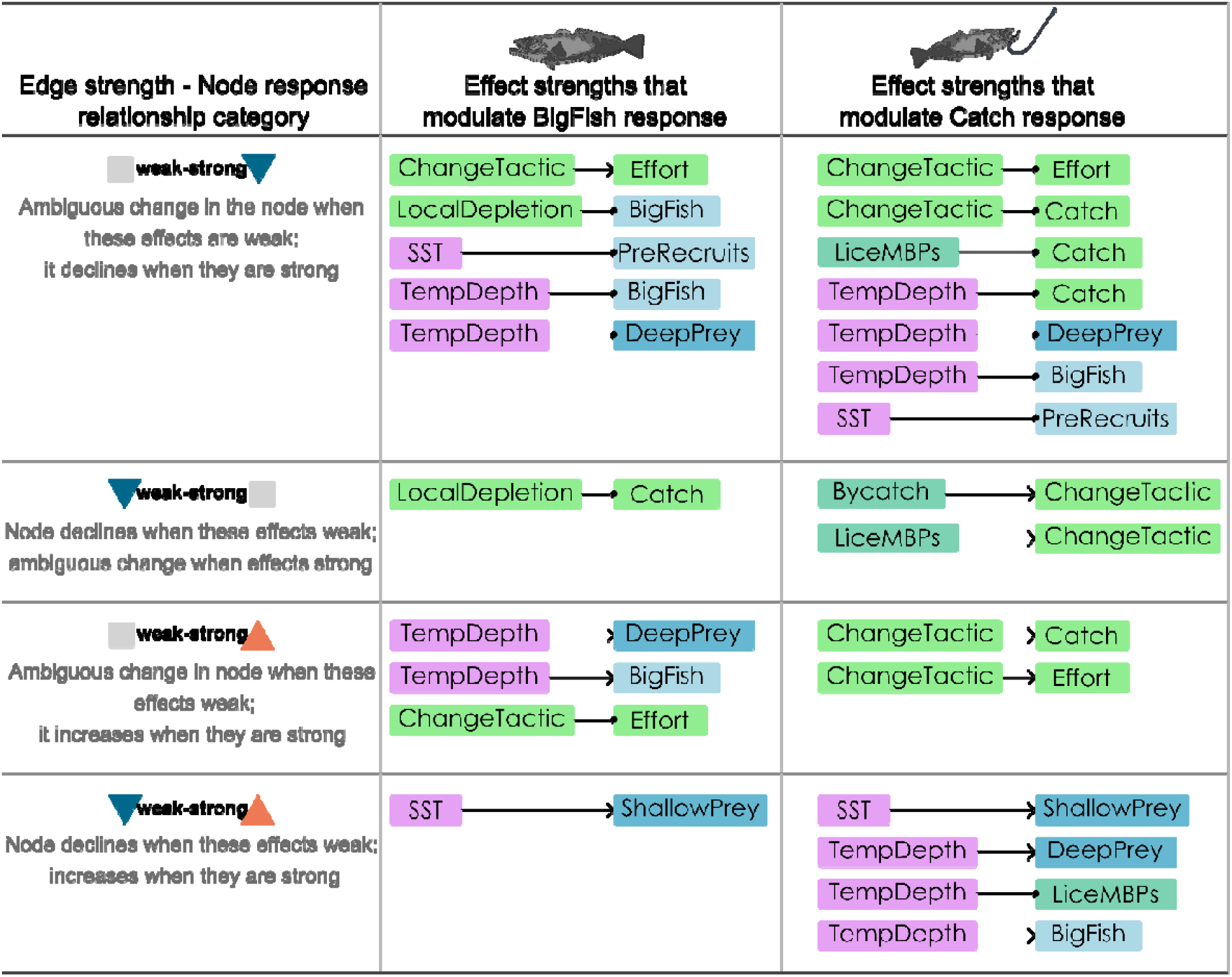
Categories of effect strength – node response relationships. For scenarios where BigFish and Catch qualitative responses (sign) were sensitive to the strength of direct outbound effects from the perturbed node, the nature (shape) of the relationship is summarised into categories. The observed categories are described in column 1. Columns 2 and 3 list the effect strengths (originating from the node being perturbed) to which BigFish and Catch node responses are sensitive as described for each category.

### Overall results

Results from each of our three sets of analyses provide different information about the likely responses of system components to press increases in each of the drivers. Table 3 synthesises the findings for each driver.

**Table 3:** Synthesis of results for each driver from across the 3 different analyses.

| Driver<br>(press<br>increase) | Synthesis of findings |
| --- | --- |
| SST | Mean changes across model versions suggested most variables declined in response to increased SST (Fig. 2), but subsequent analyses show those predicted changes are less clear cut. Changes in toothfish and catch were ambiguous when higher SST positively affected shallow prey (Fig. 3), because the strength of that effect modulated the direction (sign) of change in catch (Fig. 5a). |
| Temp. at<br>depth | Averaged across all model versions, responses to increased TempDepth were mostly ambiguous (Fig. 2) due to the number of unknown effects and sensitivity to both the sign and strength of TempDepth effects. Catch was most likely to decline if LiceMBPs were positively affected by TempDepth, but sometimes increased if LiceMBPs strongly negatively impacted Catch and DeepPrey strongly benefitted from higher temperatures (Fig. 3, Fig. 6d, Supp. Fig S3). BigFish were most likely to decline if they and DeepPrey were strongly negatively affected by TempDepth, but could increase if those nodes benefitted from warmer water (Fig. 3, Fig. 6a, SFig. S3). |
| Down-<br>welling | Average responses to increased downwelling were ambiguous (Fig. 2). The main mechanism by which downwelling appeared to affect the system was via its contribution to deep water temperature, rather than on shallow nodes. Flow-on effects from TempDepth are likely to determine the nature of the responses (Figs. 3, 4, S2). |
| Change<br>Tactic | The impacts of changing fishing tactic depended on how it affected effort and catch (Fig. 3, 5c). More detailed representation of fishery behaviour within the model is required for a more complete understanding of its impact on outcomes. |
| Local<br>Depletion | Greater localised depletion caused most nodes to decline (Fig. 2), but when the effect of depletion on toothfish was weak, or the effect on catch was strong, then toothfish decline was less certain (Fig. 5d). |
| Bycatch | Increased bycatch caused most variables to decline (Fig. 2), but if the fishery is responsive to the change, then it may be able to reduce the certainty of those declines (Figs. 3 & 5e). |
| LiceMBPs | Average responses to increased LiceMBPs were ambiguous (Fig. 2), due to sensitivity to the strength of their effect on ChangeTactic (Fig. 5b), and on the sign of ChangeTactic effects on Effort and Catch (Fig. 3). Catch and Landings tended to decrease if the effect on ChangeTactic was weak, and/or effect on catch was strong, and/or if ChangeTactic negatively affected Effort or Catch. BigFish could potentially increase in those scenarios, but largely had an ambiguous response. |
| Depredation | Landings declined in response to greater depredation (Fig. 2) and subsequent analyses show that this result was not sensitive to the sign or strength of effects - the presence of the effect is enough to cause certain decline (Fig. 5f). |

## Discussion

We have produced the first qualitative network representation of the HIMI toothfish population, fishery, and associated ecosystem, and have used this model to predict how the system is likely to respond to environmental and fishery changes. We found key system components (toothfish, catch, landings) had the potential to decline in response to press increases of both environmental and fishery drivers. Our approach for capturing uncertainties and unknown/ambiguous effects within a set of alternative model versions enabled us to investigate when and how the different mechanisms of change are important, and look at key interactions determining outcomes (a visual summary is provided in Figure 7). Responses to environmental drivers varied depending on how increased temperature affected different parts of the system, highlighting the importance of addressing knowledge gaps in that area. Responses to fisheries drivers tended to be sensitive to the way fishery behaviour affected effort and catch, indicating the potential for the fishery to mitigate negative impacts of those drivers. The qualitative responses also depended on the magnitude of perturbation effects on different components of the system, and in some cases, effect strength modulated the likely sign of node responses. For example, the magnitudes of temperature effects on shallow prey and on benthic predators (such as sea lice) determine whether catch is more likely to increase or decrease.

**Figure 7:**
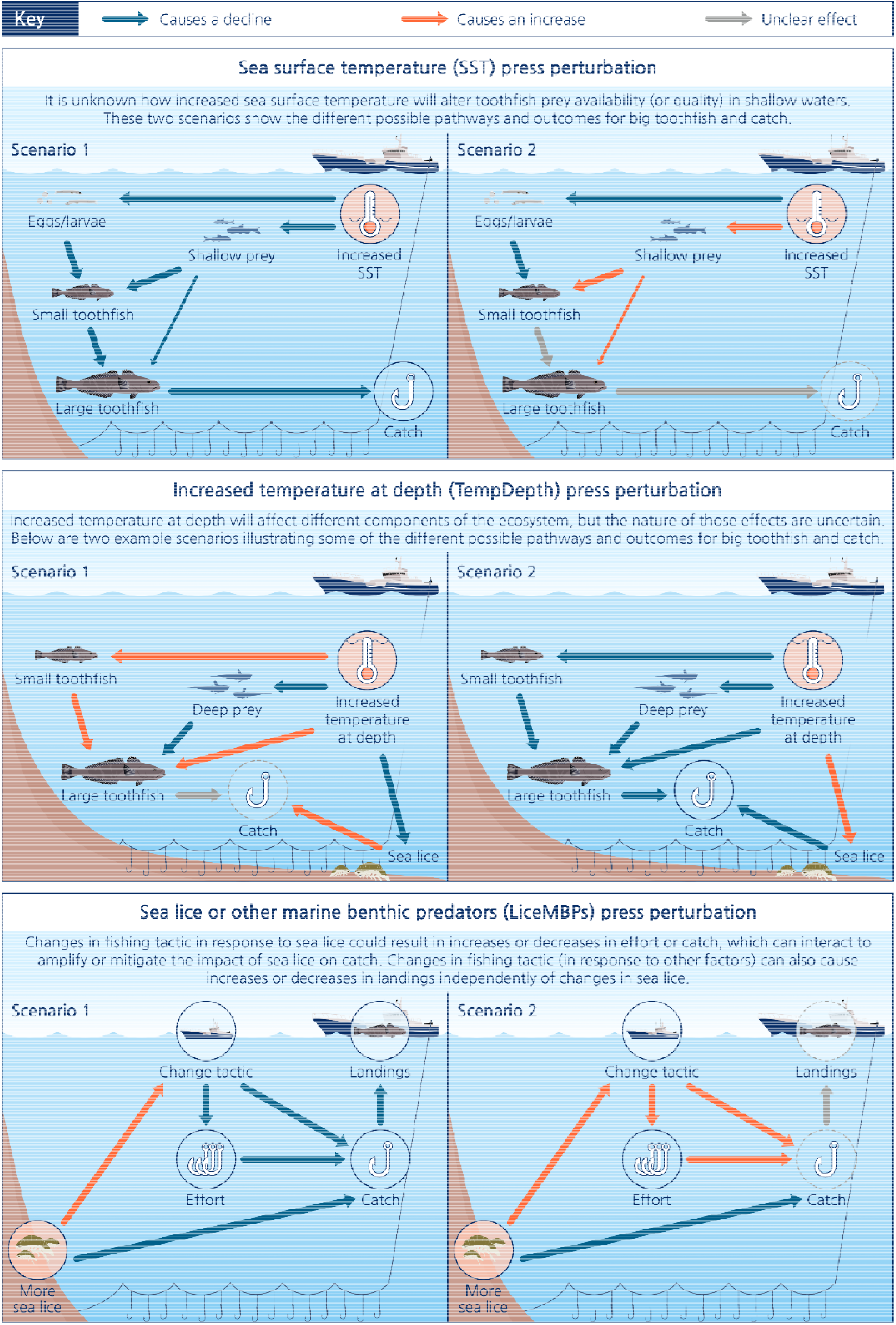
infographic illustrating different scenarios for test cases where responses are conditional on the sign of unknown edges. Note that when a node is being affected both positively and negatively by a perturbation, the relative strength of those effects is important in determining the direction of change in that node, e.g. in the top-right panel, the indirect effects of increased SST on SmallFish are opposing.

This work has highlighted the importance of the unknown effects of temperature on the responses of the system to environmental drivers. Increasing SST caused declines in toothfish population and catch when the shallow-water prey-field was negatively affected by warmer water, but when the toothfish prey-field improved, then the sign of their responses depended on the magnitude of that effect. This is consistent with other modelling and observational studies that have found warmer SST correlates with reduced productivity at lower trophic levels, and longer, less efficient food chains in this region (e.g. Venkataramana et al. 2019; Subramaniam, Melbourne-Thomas, et al. 2020). In this scenario, we would expect a lag between the increase in SST and subsequent impact on toothfish numbers at depth, i.e. the generation time for smaller toothfish directly affected by prey-field changes to mature and migrate deeper. Warm SST anomalies can indicate a shift away from the upwelling regime to which local species are adapted (Gille et al. 2014), and though we did not assume a resultant shift in prey species composition would be detrimental for toothfish, our results indicate a sensitivity to both the sign and strength of that effect.

Downwelling, in addition to diminishing productivity, is also a mechanism for transporting warm water to depth (Su et al. 2021). We found that qualitative changes in response to increased temperature at depth (either direct increase of TempDepth or via downwelling) depended on how warmer temperature affected different components of the system. Specifically, toothfish population change was sensitive to the effect of higher temperature on deep prey as well as direct effects on toothfish, but catch was most sensitive to the effect of temperature on sea lice. These predicted responses could either be immediate (e.g. due to prey distribution shifts, or sea lice), or involve a lag (e.g. via altered prey-field quality). No studies in this region have investigated potential ecological implications of warming water at depth, and our findings highlight the need for further research to clarify how the warming trend being observed in the upper 2000 m of the Southern Ocean will affect this fishery (Gille 2002; Meredith et al. 2019). Nonetheless, the circumpolar distribution of Patagonian toothfish includes water temperatures ranging 2-11°C as compared to about 2°C around HIMI (van Wijk et al. 2010; Collins et al. 2010; Mintenbeck 2017), so it is likely that indirect effects on toothfish via their prey, or on catch via sea lice, will be more important than direct effects.

Responses to changes in the fishery or fishery-ecosystem interactions were varied. Greater localised depletion of toothfish by the fishery and increased bycatch both caused declines in toothfish population and catch, and increased depredation caused decline in landings. However, while our results suggest fairly certain declines in landings in response to these drivers, we cannot comment on the magnitude of those declines using a qualitative model. Quantitative ecosystem models of the Kerguelen Plateau indicate that fishing is not a major driver in the system (Subramaniam, Corney, et al. 2020; Subramaniam, Melbourne-Thomas, et al. 2020) and depredation, while it causes substantial loss of catch, especially in the French toothfish fisheries to the north (Tixier et al. 2020), is not yet of major concern in the HIMI fishery.

### Key knowledge gaps

Our approach has enabled identification of key data gaps that must be targeted to better predict how this system responds to change, and the sustainability of the fishery into the future. A fundamental knowledge gap is how toothfish will respond to increasing water temperatures, either directly via their own thermal sensitivity and or indirectly via likely changes to their prey-field (across their depth gradient). Addressing the latter would entail predicting changes in species composition and abundance, as well as implications of those changes for toothfish (e.g. catchability/energetic cost to capture vs energetic value of prey species). Adult toothfish are opportunistic consumers (Collins et al. 2010), but behavioural and metabolic changes in both prey and predator could interact to make energetic consequences for toothfish difficult to predict. Our findings indicate that addressing this gap will be important for improving predictions of change in this system.

An unexpected result was the sensitivity of toothfish catch to the representation of sea lice (or other benthic predators) in the model. Sea lice were included in the model based on input from fishers who indicated that sea lice are a growing concern. The model sensitivity to sea lice highlights the lack of knowledge about the ecology of these organisms (Kaiser et al. 2007; Kaiser 2014). Scavenging sea lice (Order Isopoda) and other benthic predators have the potential to greatly diminish catch if their abundance increases because they remove bait from hooks. In our model, sea lice affect catch but do not interact directly with any other components of the ecosystem; however alternative network structures are possible. For example, potential interactions between toothfish and sea lice or other scavengers could include predation (e.g. Garcia de la Rosa et al. 1997), active avoidance (e.g. Yau et al. 2002), or competition for carrion, in which case lice and other scavengers could benefit from reduced toothfish numbers. Feedbacks created by such interactions, if they exist, would alter the expected responses to perturbation from those observed here. Refining representation of sea lice and other scavengers in the model, including their ecology and thermal sensitivity, will be important for clarifying predictions and informing longer term fishery planning.

### Contribution to QNM from a methodological perspective

We have presented a novel approach for exploring mechanisms by which changes flow through a network to affect responses of key nodes. Some changes, such as increased temperature at depth, are likely to affect many components of the ecosystem at the same time, and determining which effects are most important for the nodes of interest is important but challenging. QPress simulates quantitative community matrices with different combinations of edge strengths (Melbourne-Thomas et al. 2012; Ward et al. 2022). Our approach used that information to investigate how responses to press perturbation vary according to the strength of the outbound edges from the node being perturbed. The resultant plots enable comparison of the relative influence of the different pathways of perturbation impacts on key variables. For example, increased bycatch (e.g. of toothfish prey) tended to cause declines in toothfish and catch, but Figure 5e illustrates the potential for the fishery to modulate the net effect on catch through the strength of their behavioural response to increased bycatch.

Our approach has also enabled a more nuanced understanding of the interplay between edge sign (for those edges where it is unknown) and edge strength in shaping responses of nodes to press perturbation, as compared to the coarser analysis of mean responses across simulations. For example, Figure 3 shows that the direction of change in catch in response to increased temperature at depth is sensitive to the effect sign of temperature on sea lice. However, Figure 6e provides more nuanced insight into the likely responses when that effect is negative vs positive – namely, that when higher temperature negatively affected sea lice then catch tended to increase when the effect is strong and decline when it is weak, but when higher temperature positively affected sea lice, then catch consistently declined. Similar conditional information could be determined from symbolic analysis of a community matrix for smaller networks (up to 5 nodes), but can be computationally unfeasible for larger networks such as this one (Dambacher et al. 2003). This new approach therefore makes a useful progression in methodological capacity for QNM and we recommend its uptake for exploring conditional and interacting responses to perturbation in QNMs.

Given the number of unknown effects (and therefore model versions) and perturbation scenarios, we have largely opted to restrict our analyses to a single level of complexity. For example, we investigated only a single level of the mechanism pathways (i.e. the direct effects from the node being perturbed) but in some cases (e.g. downwelling perturbation) the strength (and sign) of flow-on effects are also likely to be important. The importance of these flow-on effects could be explored using the same approach noted above. In the supplementary information we also further discuss our approach for handling the many unknowns edge signs in the model.

### QNM as a tool to explore drivers through knowledge co-production

Transdisciplinary knowledge co-production entails bringing together researchers (from different disciplines) and people from other sectors (in this case, industry and resource managers) to generate knowledge that can assist in decision-making (Norström et al. 2020). The nature of the question we were attempting to address – regarding complex interactions between the biophysical effects of environmental change and fishery responses – required not only an interdisciplinary approach but also specific industry knowledge regarding observed change and industry responses. Qualitative network models proved to be an effective tool for developing a shared understanding of the system, and for refining this understanding through an iterative process (typical of knowledge co-production; Norström et al. 2020). Underpinning the model development and analysis were other communication approaches such as workshops and the use of visual summaries of model outputs (such as infographics, e.g. Figure 7). Overall, this process enabled new insights about the sensitivity of the HIMI ecosystem and fishery to environmental change – as well as the identification of key knowledge gaps – that would not have been possible without an integrated research approach that brought together expert knowledge from both research and industry perspectives.

Use of such approaches are increasing, and qualitative network models have been used in a range of other contexts to elicit and consolidate various types of expert knowledge, including as a precursor to the development of more complex, quantitative models of socioecological systems (e.g. Fulton et al. 2015). To our knowledge this is the first documented use of such an approach for a coupled biophysical-fishery system for the Southern Ocean. Future use of knowledge co-production for Southern Ocean ecosystems and fisheries could be used to support ecosystem-based management approaches and outcomes (see also Solomonsz et al. 2021).

### Next steps

We found multiple potential pathways of decline for toothfish population and catch, but also some possible mechanisms of increase (and potential for fishery to mitigate declines) that are dependent on magnitude of certain effects. The next steps must be to close key data gaps identified through this study, including sea lice ecology, thermal tolerances of different species, and the changes in toothfish prey-field in relation to oceanographic changes. Until now, information about sea lice interactions with the fishery has been mostly anecdotal, with some data collected informally; however, in response to this study the industry has commenced collecting data and samples more formally, with the aim of better understanding where and under what conditions sea lice occur. It will also be important to hone understanding of the likely strength of effects (and thus most likely outcomes) for the fishery to plan its next steps.

As these key data gaps are filled, the findings can be integrated into single species and ecosystem models that spatially and temporally resolve key patterns and drivers of the Kerguelen Plateau (e.g. Subramaniam, Melbourne-Thomas, et al. 2020; Subramaniam et al. 2022) to inform fisheries management and planning using quantitative projections. Qualitative network models represent a specific timeframe (in this case multiple years to decades); however fisheries planning occurs for multiple time scales, including day-to-day decisions, seasonal plans and longer term strategies and investment decisions. With improved understanding of the effects of climate change on toothfish and the Kerguelen Plateau ecosystem, tools such as marine heatwave forecasts (Hobday et al. 2016) could help to inform shorter-term tactical decision-making for the region. Additionally, our QNM model can be updated as current uncertainties in the network are resolved to help inform system-level predictions for future change and longer-term planning and adaptation approaches.

From an industry perspective this study has unpacked and shed light on some of the potential pathways that may be seen on the target species and broader ecosystem moving forward in the HIMI fishery. This work will help focus research and operational efforts in the short to medium term which will hopefully allow for more successful long-term planning of assets and strategy.

## Supplementary information and data availability

Supplementary methodological details, results and discussion are available in the supplementary information. Additionally, R codes and simulated data are provided in the online supplementary materials.

## Author Contributions

Conceptualization: All authors

Model development and validation: DWa, JMT, RA, MM, DWe, PZ, NH, SC

Method development: DWa, SW, JMT, SC, NH

Coding and implementation: DWa

Analysis: DWa with input from all co-authors

Funding acquisition: SC, NH, JMT, DWe, PZ

Visualization: DWa

Writing – original draft: DWa

Writing – review & editing: DWa, JMT, SC, NH, DWe, RA

## Acknowledgements

This work was funded through the Australian government’s Fisheries Research Development Corporation grant FRDC2018-133. We would like to thank personnel from Austral Fisheries, Australian Longline, the Australian Fisheries Management Authority and the Australian Antarctic Division for their engagement in and support of this work, including attendance at workshops and contributing to the development of the qualitative network model. We acknowledge that the majority of this work was conducted on the traditional country of the Muwinina people, and recognise the Tasmanian Aboriginal community as custodians of land- and sea-Country.

