## Supplementary_Material for "Exploring mechanisms of change in a Southern Ocean fishery with a co-produced network model"

**Supplementary materials for “Exploring mechanisms of change in a Southern Ocean fishery with a co-produced network model”**

Delphi Ward, Nicole Hill, Jess Melbourne-Thomas, Dirk Welsford, Rhys Arangio, Malcolm McNeill, Simon Wotherspoon, Philippe Ziegler, Stuart Corney

*** Example code and data located in additional supplementary materials**

### Supplementary methods details:

#### QNMs

Analysis of QNMs is based on the community matrix (A), the elements of which are the pairwise interaction coefficients (a_ij_) representing the effect of node *j* on node *i*. The net effects of each node on each of the other nodes is captured in the adjoint of the community matrix, adj(-A). The negative inverse community matrix (-A^-1^ = adj(-A)/ det.(-A)), provides information on how nodes will respond to the press perturbation of another node (Nakajima 1992; Dambacher, Li and Rossignol 2002). Specifically, elements of the matrix -A^-1^ indicate how abundances or densities of each node would differ in a new equilibrium state, compared to the initial equilibrium state, following a press increase of another node (i.e. they do not indicate trends).

#### Assumptions in the HIMI Patagonian toothfish fishery-ecosystem QNM

Key assumptions in developing and refining the model include: 1) that the fishery is (and will continue to be) well managed, and that quota is never at a level that results in irreversible change to the toothfish population. This assumption is supported by other studies (Brooks, Epstein and Ban 2019; Ziegler and Welsford 2019; MSC 2020). In this context, catch does have a negative effect on toothfish, but it is uncertain (edge only present in 50% of simulations). 2) That high SST results in reduced recruitment of subsequent age classes. This is supported by evidence from another sub-Antarctic location (Belchier and Collins 2008). 3) That higher temperatures affect toothfish catch via changes in toothfish distribution or behaviour making them less susceptible to being caught. This assumption is based on expert advice from captains who operate fishing boats at HIMI, but is uncertain, and so the effect (where TempDepth suppresses the BigFish•-> Catch edge) is only present in 50% of simulations for each model. 4) We assumed increased water temperature would directly affect multiple ecosystem components, but made no assumptions with regards to the nature (sign) of those effects, as described in the main text. These and other minor assumptions are detailed in the supplementary table (STable S1).

#### Model testing and validation:

To test and validate the model, the network was divided into two sub-models: a biophysical sub-model and a fishery sub-model. A set of test cases based on expected system behaviour was developed to test these two sub-models separately. Specifically, the test condition we set for the biophysical sub-model was that a decrease in spawning/recruitment should result in an eventual decrease in large toothfish. The test conditions we set for the fishery sub-model were that a) an increase in toothfish stock should result in an increase in catch (via increased quota); b) higher quota should result in higher catch and landings (but at lower equilibrium stock size, given the assumption above that the fishery continues to be sustainably managed); and c) greater localised depletion should reduce stock, catch and landings. The model was refined based on this testing and continued consultation with industry.

### Table S1: Summary descriptions of nodes and their associated edges

|  | **Node** | **Description** | **Edges (outbound or inbound)** |
| --- | --- | --- | --- |
| **Toothfish lifestages** | 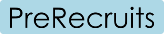 | This node captures all stages from spawning to recruitment to the juvenile population, including pelagic egg and larval stages. | - Input from BigFish via spawning. - Can be predated by ShallowPrey. - Negatively impacted by downwelling and higher SST (e.g. Belchier & Collins 2008). - Contribute to SmallFish population |
|  | 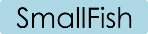 | Node captures toothfish from when they are recruited into the juvenile population until they reach a size (depth) at which they are targeted by the fishery. | - Recruited from PreRecruit population. - Consume ShallowPrey. - Can be cannibalised by BigFish (50% of sims). - (Other predation contained within the negative self-effects – not explicitly included to simplify the model) - Unknown effect from increased temperature at depth. - Contribute to BigFish population. |
|  | 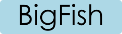 | This node represents all toothfish that are deep enough to be captured by the longline fishery, and as such will include both immature and mature individuals above a certain size. Adult toothfish are opportunistic predators (eat anything they can) and have greater natural buoyancy than juveniles. Large adults inhabit waters ranging 500-2000m+, but mostly occur in deeper waters (1200-2500m) (Péron *et al.* 2016; Farmer *et al.* 2019). They annually migrate to specific locations in shallower waters to spawn, but do skip in some years (Belchier and Collins 2008; Boucher 2018). | - Recruited from SmallFish population. - Consume DeepPrey. - Can consume ShallowPrey (50% of sims). - Unknown effect from increased TempDepth. - Consumed by DeepPredators. - Reduced by Catch (50% of sims – assumed to be well-managed so quota never set beyond what is sustainable). - Reduced by increased localised depletion (only in that test case). |
| **Toothfish-related ecosystem** components | 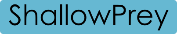 | This node represents the range of prey consumed by small toothfish. Juvenile toothfish forage above the seabed on pelagic and semi-pelagic species including fishes, euphausids, amphipods (Collins et al. 2010). An increase or decrease in this node could represent changes in numbers, but could equally represent a shift in the prey field towards better/worse quality prey for toothfish. | - Can consume PreRecruits (50% of sims) - Unknown effect of increased SST (could increase or decrease prey availability and/or quality) - Negatively affected by downwelling. - Consumed by SmallFish - Can be consumed by BigFish (50% of sims) |
|  | 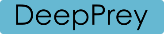 | Represents the range of organisms consumed by large toothfish at depth. This includes demersal and pelagic fish, crustaceans, cephalopods and carrion (Collins et al. 2010). As with ShallowPrey, an increase or decrease in this node could represent shifts in the quantity and/or quality of prey available to toothfish. (Equally, a lack of change could still involve a shift in the preyfield, but one that does not entail net change in the availability of energy/nutrients for toothfish). | - Negatively affected by bycatch - Unknown effect from increased temperature at depth (could increase or decrease prey availability and/or quality) - Consumed by BigFish |
|  | 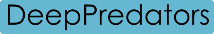 | Represents predators of BigFish. This is maintained as separate node (unlike shallow predators) because sperm whales, a deep-diving predator, is also the main depredating species in the HIMI fishery. Other deep-living or deep-diving predators include: elephant seals, giant squid and sleeper sharks (Collins *et al.* 2010). | - Consume BigFish - Contribute to depredation (50% of sims) |
| **Fishery-related ecosystem components** | 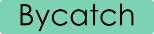 | Bycatch species vary spatially, but the main by-catch are macrourids and skates (both subject to daily limits and seasonal quotas). Macrourids are important toothfish prey, and their swim bladder prevents them being released once caught (due to ‘decompression trauma’). | - Increase with increased fishing effort (50% of sims – subject to quota) - Triggers ChangeTactic (50% of sims – when daily limits or seasonal quotas are being approached) - Negative effect on DeepPrey. |
|  | 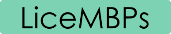 | Sea lice and other marine benthic predators. These can remove bait from hooks which limits toothfish catch. They can also nibble toothfish caught on the lines, but this is of less concern. Sea lice are crustaceans belonging to Order Isopoda, which is a diverse order that includes genera containing free-living, scavenging species (Kaiser, Barnes and Brandt 2007; Kaiser 2014). Little is known about their ecology or thermal tolerances. | - Unknown effect from temperature at depth. - Negative effect on Catch. - Causes change in fishing tactic (ChangeTactic) e.g. fishing boats move away from area of lice attack. |
|  | 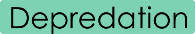 | In the HIMI fishery, depredation is mostly by sperm whales, in some months when 2-5% of lines can be affected. Not currently of major concern, but it is important in the French Kerguelen fishery to the north. | - Positive effect from DeepPredators (50% of sims) - Negative effect on Landings |
| **Fishery** | 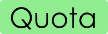 | This represents fishery-management processes and decisions to ensure sustainability of the fishery. Quota is set based on toothfish population estimated by surveys, tag-recapture data, modelling and expert input. | - Positively affected by BigFish population (a population growth would mean higher possible quota). - Positively affects Effort. |
|  | 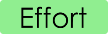 | Represents fishing effort, and includes soak time, number of hooks set and bait used. | - Positively affected by Quota. - Unknown effect from ChangeTactic. - Positive, uncertain effect on Bycatch (50% of sims). - Positive effect on Catch. |
|  | 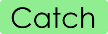 | Represents toothfish caught on hooks (which may be subsequently brought onboard as Landings or depredated by sperm whales). | - Positively affected by Effort - Positively affected by BigFish - Negatively affected by LiceMBPs (they remove bait from hooks). - Negatively affected by LocalDepletion (can’t catch fish where they have been depleted) - Unknown effect from ChangeTactic. - Negative, uncertain (50% of sims) effect from TempDepth – arising out of the hypothesized mediating effect of temperature on toothfish distribution or behaviour making them less ‘catchable’. - Positively affects Landings. - Negatively affects BigFish (50% of sims – assumption of sustainability). |
|  | 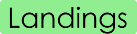 | This represents Catch that has been successfully brought onboard, i.e. not depredated. | - Positive input from Catch - Negative effect from Depredation. |
|  | 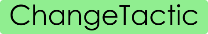 | This node attempts to capture aspects of fisheries behaviour that could affect effort and catch. Although it is not specific, this could include for example, changing fishing locations (e.g. in response to sea lice or weather conditions), changing bait used or fishing depth. | - Positively affected by LiceMBPs - Positively affected by Bycatch (50% of sims – e.g. once bycatch limits approached). - Negative effect from Landings (50% of sims – less likely to change tactic if landings are increasing). - Unknown effect on effort (could cause increase or decrease). - Unknown effect on Catch (could cause increase or decrease). |
|  | 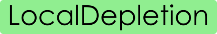 | This node represents the scenario where localised depletion of toothfish occurs. As a ‘hand-of-god’ node, it only affects the network when it is directly perturbed. | - Negatively affects BigFish. - Negatively affects Catch. |
| **Environmental pressures** | 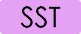 | Sea surface temperature. As a ‘hand-of-god’ node, it only affects the network when it is directly perturbed (i.e. increased). | - Negatively affects PreRecruits (Belchier & Collins 2008). - Unknown effect on ShallowPrey (could increase or decrease quantity and/or quantity of shallow prey). |
|  | 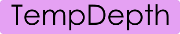 | Temperature at depth. This node only affects the network when it or Downwelling are directly perturbed. It is largely unknown how increased temperature depth will affect different components of the system. | - Unknown effect on DeepPrey - Unknown effect on SmallFish - Unknown effect on BigFish - Negative, uncertain (50% of sims) effect on Catch (arising from mediated effect on toothfish ‘catchability’). - Unknown effect on LiceMBPs |
|  | 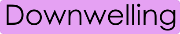 | The system is largely an upwelling system, but shifts in winds can cause downwelling, which could transfer heat to deeper waters. As a ‘hand-of-god’ node, it only affects the network when it is directly perturbed. | - Negative effect on PreRecruits - Negative effect on ShallowPrey - Positive effect on TempDepth |

### Supplementary results

#### Figure S1

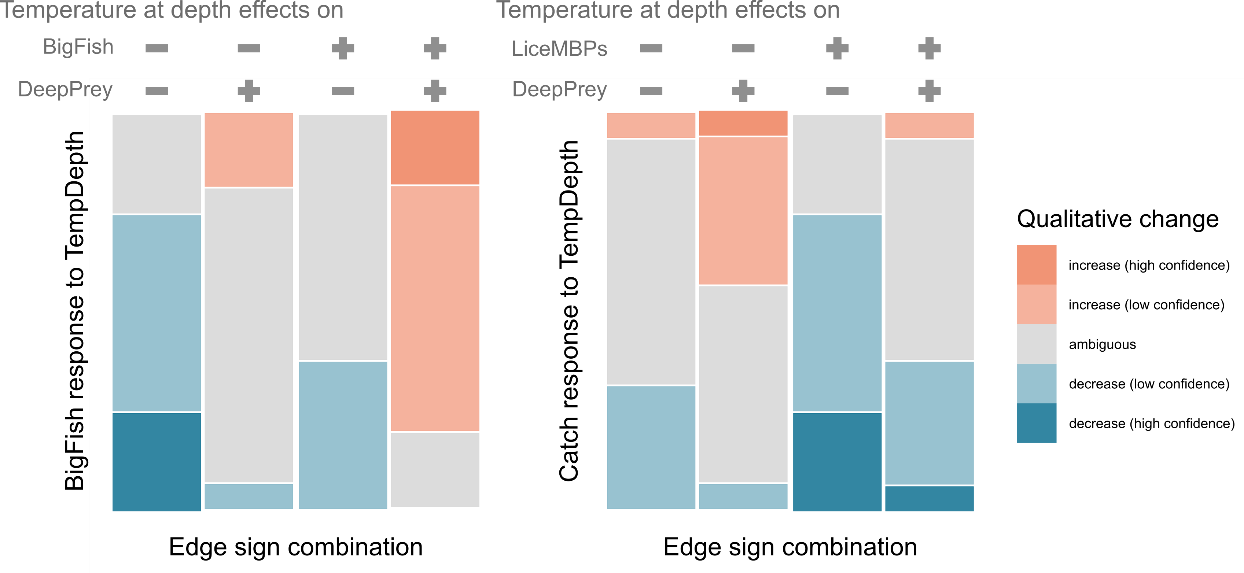

**SI Figure 1: Mosaic plots illustrating how responses to increased TempDepth vary according to sign of temperature effects.** The plot on the left shows BigFish responses in models grouped by the sign of TempDepth effect on BigFish and on Deep Prey (the sign of these two edges explain the most variability in BigFish response). The plot on the right displays the responses of Catch in model versions grouped by the sign of the effects of TempDepth on Lice and other marine benthic predators, and on Deep Prey of toothfish (the sign of these two edges explain the most variability in responses of Catch and Landings). Each mosaic plot represents the responses of nodes across all 128 model versions, and each column represents the responses in a different subset (n=32) of those.

#### Figure S2

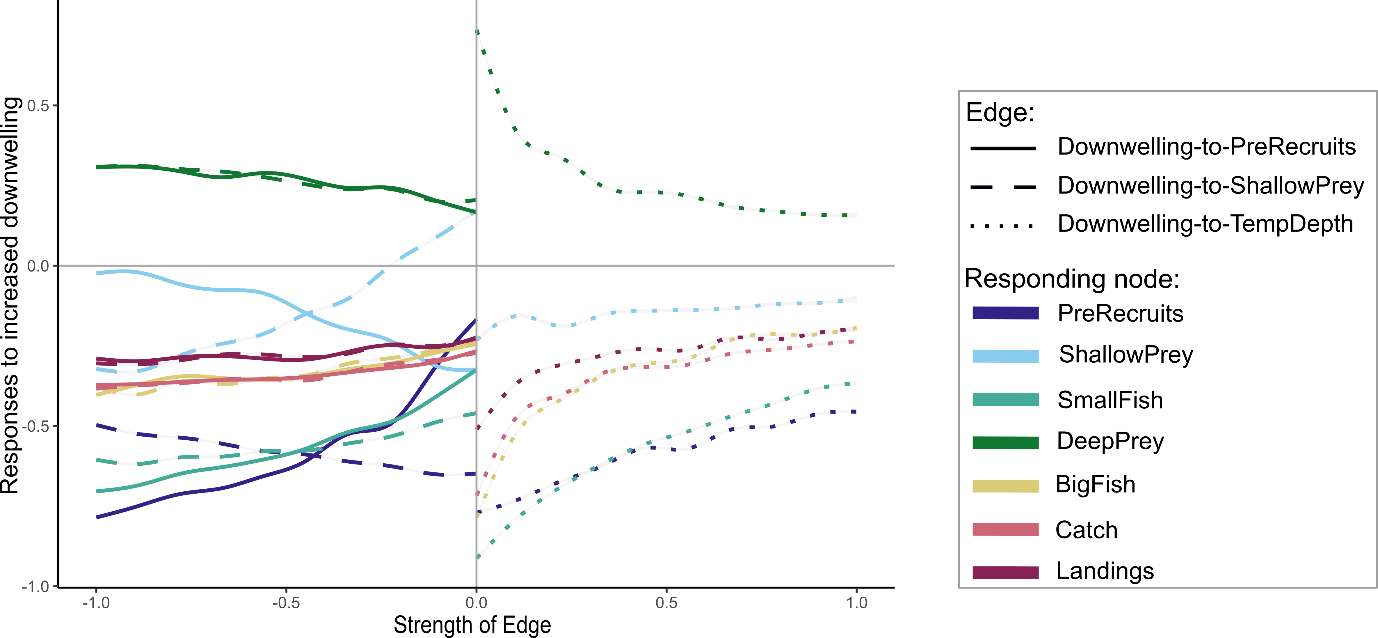

**SI Figure 2: Relationships between qualitative responses to downwelling perturbation and effect strengths.** Each line is a gam smooth of the qualitative responses of one node against one of the outbound edges downwelling. Colours correspond to the response of one node to downwelling perturbation. Line styles correspond to the different edges, and values on the x-axis correspond to the strength of the edges. Values on the y-axis correspond to the qualitative responses of the nodes, with -1 indicating a decline, +1 an increase, and 0 indicates no change (values between |0 and 1| suggest a mix of qualitative responses).

#### Figure S3

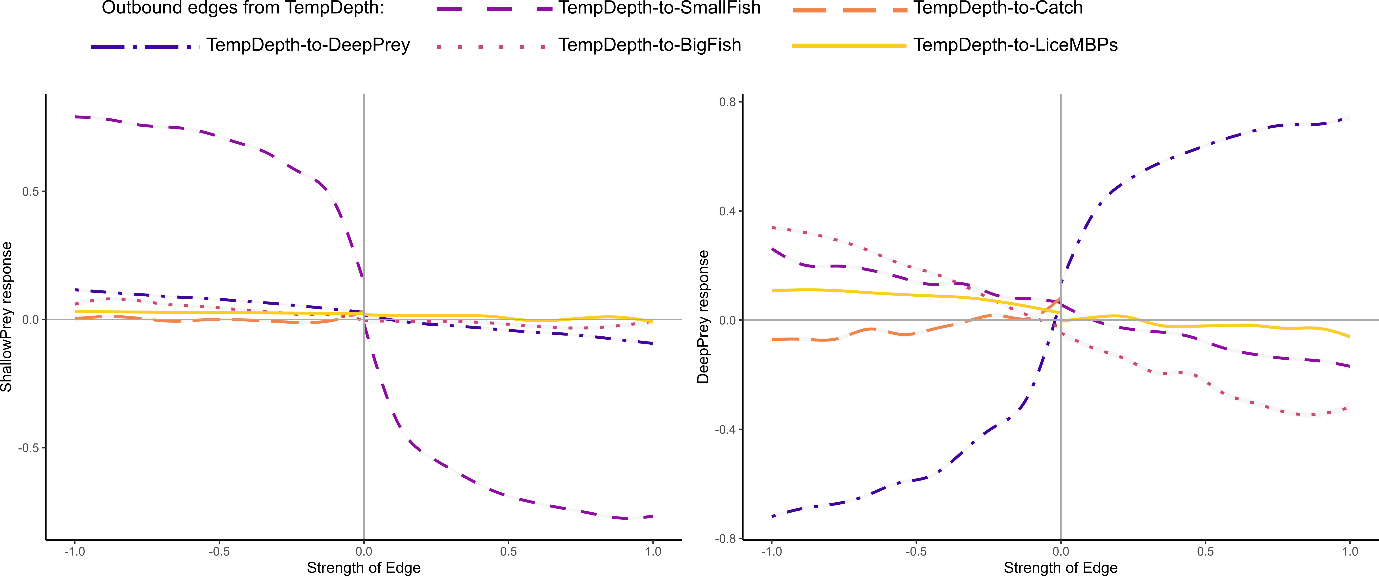

**SI Figure 3: Relationships between qualitative responses of prey to increased TempDepth according to effect strengths.** Each plot shows the responses of one node, ShallowPrey (left) and DeepPrey (right), with lines representing the relationship between the qualitative response of that node, and the strength of each of the outbound effects from TempDepth. Values on the x-axis correspond to the strength of the outward effects from the node being perturbed. Values on the y-axis correspond to the qualitative responses of the nodes, with -1 indicating a decline, +1 an increase, and 0 indicates no change (values between |0 and 1| suggest a mix of qualitative responses). Equivalent plots of BigFish, SmallFish, PreRecruits, Landings and Catch node responses shown in Figure 6 of main text.

### Supplementary discussion

**Brief discussion of approach for dealing with multiple unknown edge signs**

Our study is novel in the sheer number of unknown effect signs represented in the qualitative model. To deal with uncertainty in the way that increased water temperature will affect different components of the system, we created alternative versions of the model capturing the different possible combinations of alternative effect signs. The strength of this approach is that it has enabled us to explore the conditions under which toothfish population and catch might increase or decrease, and to identify key data gaps that need to be filled to maximise our predictive capability. However, different effect sign combinations in the alternative model versions result in different emergent feedbacks, and averaging over so many model versions (effect sign combinations) can result in average responses that are quite different compared to more extreme but possible cases (e.g. where increases in temperature negatively affect everything). To investigate the importance of specific edge signs in shaping qualitative responses, we compared the changes in the two halves of the set of model versions as distinguished by the sign of a single effect, but have largely missed more detailed conditions (e.g. combinations of effect signs, but see Fig. 4 and SIFig. 1 as examples). While we did not do so, the same approach can be used to explore co-dependencies on different effects in shaping qualitative responses to perturbations, for example to explore consequences of particular management responses to perturbation (e.g. in a model where this is captured in more detail).
